# PEPTERGENT: A Peptide-Based Method for Detergent-Free Extraction and Purification of Membrane Proteins and Membrane Proteomes

**DOI:** 10.64898/2026.03.17.711971

**Authors:** Frank Antony, Ashim Bhattacharya, Franck Duong van Hoa

## Abstract

Peptergent is a novel class of amphipathic peptides that enable detergent-free extraction and purification of membrane proteins (MPs). These designed peptides self-assemble around hydrophobic transmembrane regions of proteins, forming stable, water-soluble assemblies that can be isolated directly from biological membranes. By doing so, Peptergent bypass the limitations imposed by traditional detergents, which often destabilize proteins and restrict downstream analyses. Since detergents are completely avoided, Peptergent-isolated MPs are directly amenable to structural and mass spectrometry (MS) analysis, thereby addressing their persistent underrepresentation in proteomic datasets and improving their accessibility for drug-screening strategies. Here, we describe a streamlined protocol for isolating MPs with the Peptergent PDET-1, followed by exchange into His-tagged Peptidiscs for Ni-NTA-based affinity purification. The method comprises membrane isolation, peptide preparation, protein extraction, clarification, and exchange of MPs from Peptergent to Peptidiscs. Application of this workflow yields enriched membrane proteomes compatible with downstream LC–MS/MS analysis, with improved recovery of hydrophobic and multi-pass membrane proteins.

**Key features:**

- Direct extraction and solubilization of membrane proteins in Peptergents
- Exchange into His-tagged Peptidiscs enabling affinity purification of MPs
- 100% detergent-free workflow compatible with LC–MS/MS analysis
- Applicable to cultured cells and tissue-derived membrane fractions

**In Brief:** We describe a Peptergent-based workflow for isolating membrane proteins directly from membrane preparations. Proteins are extracted with the Peptergent peptide scaffold (PDET-1) and transferred into His-tagged Peptidisc (HD-43). The water-soluble membrane proteins are enriched by Ni-NTA affinity purification and prepared for bottom-up mass spectrometry, yielding enriched membrane proteomes and dried peptide samples ready for LC–MS analysis

**Graphical Overview:** 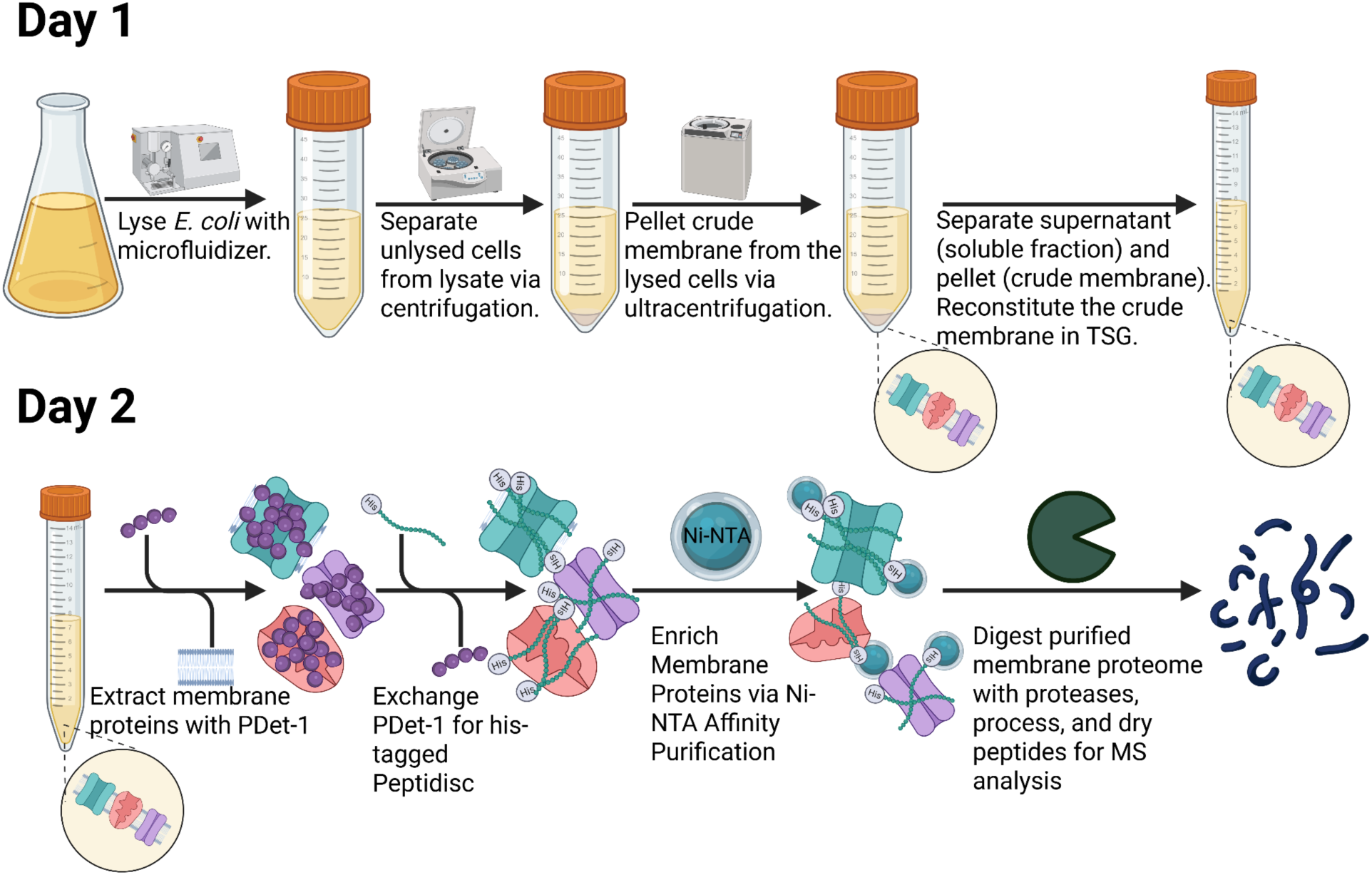

## Background

Membrane proteins (MPs) constitute ∼30% of the proteome and represent the majority of current therapeutic targets (1). However, their identification as well as their biochemical and structural characterization remains fundamentally constrained by the need to extract them from lipid bilayers while preserving native structure, oligomeric integrity, and ligand responsiveness (2–4). Conventional workflows rely on detergent solubilization followed by purification in micelles, a process that frequently destabilizes MPs, disrupts native lipid interactions, and biases downstream structural and functional analyses (3,5,6). These surfactants are also largely incompatible with downstream mass spectrometry (MS), thereby constraining quantitative membrane proteome analyses (7).

Alternative detergent-free systems, including nanodiscs and Peptidiscs scaffolds, as well as styrene–maleic acid lipid particles (SMALPs), partially address detergent-induced artifacts (8). However, scaffold-based approaches still require an initial detergent extraction, whereas SMALPs, though detergent-free, introduce charged polymers that are incompatible with MS and necessitate organic-phase cleanup steps that bias protein recovery (8,9). Consequently, a gap remains between native membrane extraction and MS-compatible proteomic analysis.

A growing body of work has demonstrated that short amphiphilic peptides can solubilize membrane proteins from lipid bilayers in a manner analogous to conventional detergents (10,11). Because these peptides combine detergent-like extraction properties with a peptide-based scaffold, they have been termed peptide detergents, or “Peptergents.” (12) Early studies showed that Peptergents efficiently solubilize membrane proteins while improving the stability and longevity of membrane enzymes compared with traditional surfactants (10,12–14, 21). Structural studies later revealed that these peptides self-assemble into ordered supramolecular architectures, including β-sheet structures that encircle hydrophobic protein surfaces and effectively replace classical detergent micelles with peptide-based amphipathic scaffolds (15,16).

Building on these observations, we describe here a simple detergent-free strategy for extracting MPs from biological membranes using amphipathic Peptergents. In this workflow, membranes are incubated with the Peptergent PDET-1, forming peptide-stabilized MPs assemblies. Because PDET-1 lacks an affinity handle, the solubilized MPs are subsequently exchanged into His-tagged Peptidiscs. This exchange step enables Ni-NTA affinity purification of reconstituted MPs depleting soluble contaminants that co-extract during membrane solubilization (17–20). We outline the key steps of this workflow, including membrane isolation, peptide preparation, membrane extraction with PDET-1, clarification, exchange into His-Peptidiscs, Ni-NTA enrichment of reconstituted MPs, and sample processing for bottom-up MS analysis. This protocol enables preparation of crude membrane fractions from cultured cells or tissues for proteomic analysis while completely avoiding the use of detergents.

## Materials and Reagents

### Biological Materials

1. Glycerol stock of Bl21 *E. coli*; Lab stock; Check Addgene (https://www.addgene.org/protocols/create-glycerol-stock/) for how to make; Store at -80°C.

### Reagents

1. PDET-1 (Peptergent); Peptidisc Biotech; Store at - 20 °C.
2. HD-43 (His-Tagged Peptidisc); Peptidisc Biotech; Store at - 20 °C.
3. Ni-NTA Agarose; Qiagen; Catalogue number 30210; Store at 4 °C.
4. cOmplete Protease Inhibitor Cocktail; Sigma-Aldrich; Catalogue number 11697498001; Store at 4 °C.
5. Mass Spectrometry Grade Proteases; Thermo Fisher Scientific; Catalogue number 90057; Store at -20 °C.
6. C18-HD Disk 47 mm; Empore (C18 Solid Phase Extraction Disk); UNSPSC code 41120000; Store at room temperature.
7. Polygoprep 300-20 C18 (C18 resin); Macherey-Nagel; Catalogue number 711025.100; Store at room temperature.
8. Tris; BioShop; Catalogue number TRS003.5; Store at room temperature.
9. Sodium Chloride (NaCl); Fisher Scientific; Catalogue number BP358-212; Store at room temperature.
10. Ethylenediaminetetraacetic acid (EDTA); BioShop; Catalogue number EDT002.500; Store at room temperature.
11. Phenylmethylsulfonyl fluoride (PMSF); Sigma Aldrich; Catalogue number P-7626; Store at room temperature.
12. Dithiothreitol (DTT); BioShop; Catalogue number DTT002.10; Store at -20 °C.
13. Iodoacetamide (IAA); BioShop; Catalogue number IOD500.5; Store at 4 °C.
14. Urea; BioShop; Catalogue number URE002.5; Store at room temperature.
15. Methanol; Sigma-Aldrich; Catalogue number 179337-4L; Store at room temperature.
16. Acetonitrile (Certified ACS); Fisher Scientific; Catalogue Number A211; Store at room temperature.
17. Imidazole; BioShop; Catalogue number IMD510.500; Store at room temperature.
18. Formic acid; Sigma-Aldrich; Catalogue number 695076-100ML; Store at room temperature.
19. Tryptone; Bioshop; Catalogue number TRP402.5; Store at room temperature.
20. Yeast Extract; Bioshop; Catalogue number YEX555.500; Store at room temperature.
21. Aluminum foil; Canada Tire; Catalogue number 053-0371-0; Store at room temperature.
22. Glycerol; Fisher Scientific; Catalogue number BP229-4; Store at room temperature.
23. Ammonium Bicarbonate; Fisher Scientific; Catalogue number A643500; Store at room temperature.
24. Sodium Hydroxide (NaOH); Fisher Scientific; Catalogue number S318-3; Store at room temperature.
25. Hydrochloric Acid (HCl) Solution; Fisher Scientific; Catalogue number A144-212; Store at room temperature.
26. Bleach; Lavo Pro 6; ULINE Canada; Catalogue number S-24408; Store at room temperature.
27. 100% Reagent Alcohol; Fisher Scientific; Catalogue number A962^F^-1Gal; Store at room temperature *This reagent is regulated and requires purchase and tracking through the university.
28. Compressed Air; Linde Canada; Please contact Linde directly to inquire how to purchase this item (1-800-225-8247).
29. Water Suitable for HPLC; Sigma-Aldrich; Catalogue number 270733-1L; Store at room temperature.
30. Protein Assay Dye Reagent Concentrate; Bio Rad; Catalogue number 5000006; Store at 4°C.

### Solutions

1. 6 M NaOH solution (see Recipes)
2. 6 M HCl solution (see Recipes)
3. 20% Ethanol solution (see Recipes)
4. 1 M Tris pH 7.8 solution (see Recipes)
5. 1 M NaCl solution (see Recipes)
6. 50% glycerol solution (see Recipes)
7. TS buffer (see Recipes)
8. 2x TS buffer (see Recipes)
9. TSG buffer (see Recipes)
10. LB media (see Recipes)
11. 50 mM ammonium bicarbonate solution (see Recipes)
12. 2 M Imidazole solution (see Recipes)
13. Ni-NTA Elution buffer (see Recipes)
14. 1 M PMSF solution (see Recipes)
15. 0.5 M EDTA pH 8.0 solution (see Recipes)
16. 100x Protease inhibitor tablet solution (see Recipes)
17. 0.5 mg/mL Mass Spec Protease solution (see Recipes)
18. 1 M DTT solution (see Recipes)
19. 500 mM IAA solution (see Recipes)
20. 2.4 mg/mL PDET-1 solution (see Recipes)
21. 5 mg/mL HD-43 solution (see Recipes)
22. Solubilization buffer (see Recipes)
23. C18 Resin Slurry (see Recipes)
24. 10% Formic acid solution (see Recipes)
25. 0.1% Formic acid solution (see Recipes)
26. 40% Acetonitrile, 0.1% Formic acid solution (see Recipes)
27. 1x Bradford Assay solution (see Recipes)

### Recipes

A useful formula for determining the amount of reagent to weigh when preparing a solution: Grams to weigh = Desired Molarity (mol/L) * Molecular weight (g/mol) * Desired Volume (L)

Use an online molarity calculator to determine adjusted values if using different quantities while preparing reagents (https://www.calculator.net/molarity-calculator.html).

1. 6 M NaOH solution (250 mL)

a. Store at room temperature

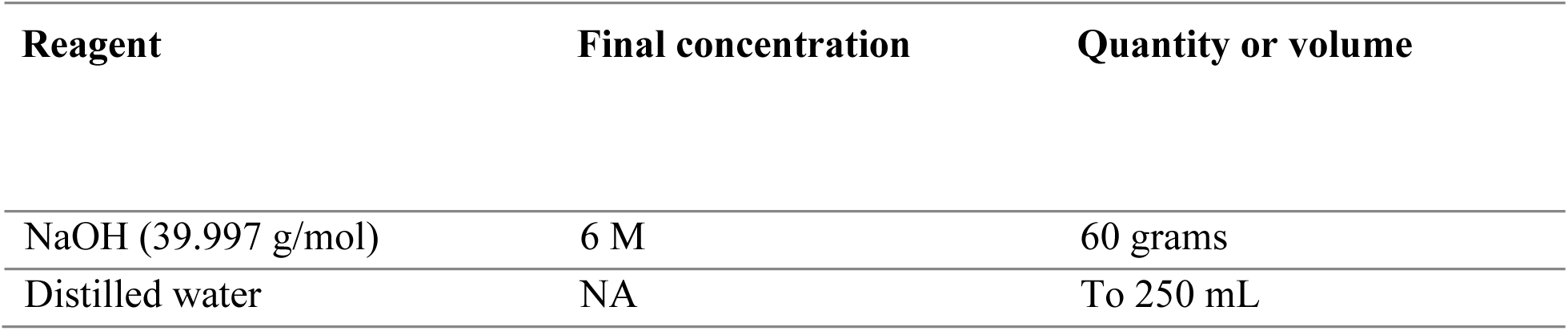

2. 6 M HCl solution (250 mL)

a. Store at room temperature

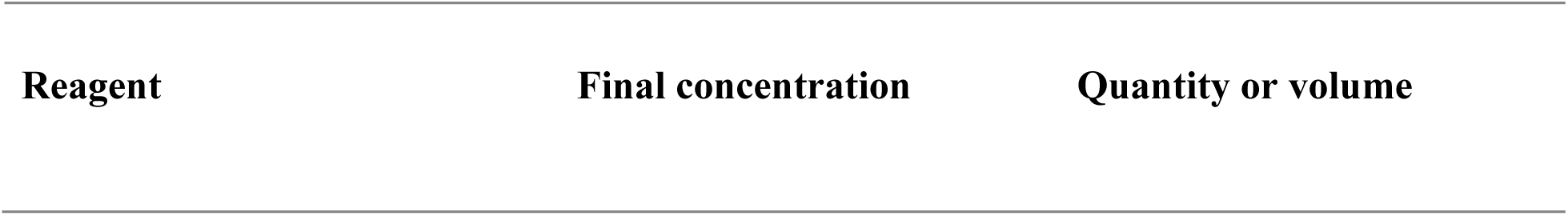
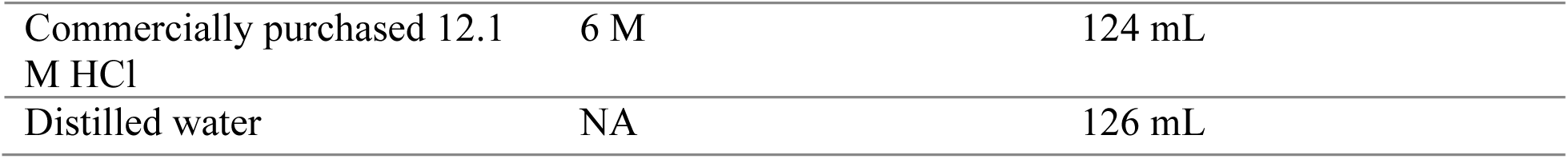

3. 20% Ethanol solution (250 mL)

a. Store at room temperature.

3. 20% Ethanol solution (250 mL) - a. Store at room temperature.
| Reagent | Final concentration | Quantity or volume |
| --- | --- | --- |
| Commercially purchased 100% Ethanol solution | 20% | 50 mL |
| Distilled water | NA | 200 mL |

4. 1 M Tris pH 7.8 solution (500 mL)

a. Store at room temperature.

4. 1 M Tris pH 7.8 solution (500 mL) - a. Store at room temperature.
| Reagent | Final concentration | Quantity or volume |
| --- | --- | --- |
| Tris (121.14 g/mol) | 1 M | 60.57 grams |
| 6 M HCl Solution | pH 7.8 | X mL to get to pH 7.8 |
| Distilled water | NA | To 500 mL |

5. 1 M NaCl solution (1 L)

a. Store at room temperature.

5. 1 M NaCl solution (1 L) - a. Store at room temperature.
| Reagent | Final concentration | Quantity or volume |
| --- | --- | --- |
| NaCl (58.44 g/mol) | 1 M | 58.44 grams |
| Distilled water | NA | To 1 L |

6. 50% glycerol solution (200 mL)

a. Avoid using a serological pipette or graduated cylinder as the glycerol will stick to the walls - more accurate to slowly pour the solution into the media bottle. Store at room temperature.

| Reagent | Final concentration | Quantity or volume |
| --- | --- | --- |
| Commercially purchased 100% glycerol solution | 50% | 100 mL |
| Distilled water | NA | 100 mL |

7. TS buffer (1 L).

a. Store at room temperature or 4 °C.

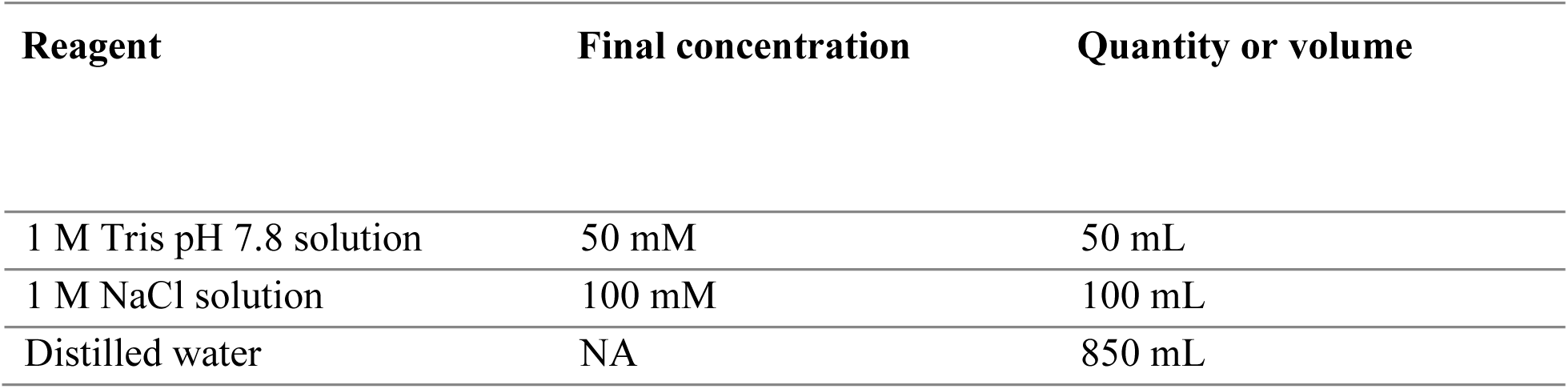

7. TS buffer (1 L). - Store at room temperature or 4 °C.
| Reagent | Final concentration | Quantity or volume |
| --- | --- | --- |
| 1 M Tris pH 7.8 solution | 50 mM | 50 mL |
| 1 M NaCl solution | 100 mM | 100 mL |
| Distilled water | NA | 850 mL |

8. 2x TS buffer (1L)

a. Store at room temperature or 4 °C.

8. 2x TS buffer (1L) - Store at room temperature or 4 °C.
| Reagent | Final concentration | Quantity or volume |
| --- | --- | --- |
| 1 M Tris pH 7.8 solution | 100 mM | 100 mL |
| 1 M NaCl solution | 200 mM | 200 mL |
| Distilled water | NA | 700 mL |

9. TSG buffer (1L).

a. Store at room temperature or 4 °C.

9. TSG buffer (1L). - Store at room temperature or 4 °C.
| Reagent | Final concentration | Quantity or volume |
| --- | --- | --- |
| 1 M Tris pH 7.8 solution | 50 mM | 50 mL |
| 1 M NaCl solution | 100 mM | 100 mL |
| 50% Glycerol solution | 10% | 200 mL |
| Distilled water | NA | 650 mL |

10. LB media (1 L)

a. Autoclave at 120 °C for 30 minutes with a small volume of water in the autoclave tray. Ensure the lid is loosely fit and autoclave tape is wrapped around the top of the bottle before autoclaving. After autoclaving, store at room temperature. If anything is seen growing inside add bleach at the volume of 10% of the combined LB media and bleach volume and mix. Sit for 30 minutes at room temperature. Then pour down the drain and chase the drain with tap water for 15 minutes.

| Reagent | Final concentration | Quantity or volume |
| --- | --- | --- |
| Tryptone (no specified molecular weight) | NA | 10 grams |
| Yeast extract (no specified molecular weight) | NA | 5 grams |
| NaCl (58.44 g/mol) | 171 mM | 10 grams |
| Distilled water | NA | To 1 L |

11. 50 mM ammonium bicarbonate (50 mL)

a. Store at 4 °C.

11. 50 mM ammonium bicarbonate (50 mL) - a. Store at 4 °C.
| Reagent | Final concentration | Quantity or volume |
| --- | --- | --- |
| Ammonium Bicarbonate (79.06 g/mol) | 50 mM | 197.65 mg |
| Distilled water | NA | Up to 50 mL |

12. 2 M Imidazole solution (50 mL)

a. Store at 4 °C.

12. 2 M Imidazole solution (50 mL) - a. Store at 4 °C.
| Reagent | Final concentration | Quantity or volume |
| --- | --- | --- |
| Imidazole (68.08 g/mol) | 2 M | 6.8 grams |
| Distilled water | NA | Up to 50 mL |

13. Ni-NTA Elution buffer (10 mL)

a. Store at 4 °C.

13. Ni-NTA Elution buffer (10 mL) - a. Store at 4 °C.
| Reagent | Final concentration | Quantity or volume |
| --- | --- | --- |
| 2 x TS | 50 mM Tris pH 7.8<br>100 mM NaCl | 5 mL |
| 2 M Imidazole solution | 600 mM imidazole | 3 mL |
| Distilled water | NA | 2 mL |

14. 1 M PMSF solution (10 mL)

a. Store at −20 °C. A precipitate may form during storage; use only the clear supernatant and avoid the insoluble fraction. Prepare fresh stock once the soluble portion is depleted.

| Reagent | Final concentration | Quantity or volume |
| --- | --- | --- |
| PMSF (174.19 g/mol) | 1 M | 1.74 grams |
| Isopropanol | NA | Up to 10 mL |

15. 0.5 M EDTA pH 8.0 solution (50 mL)

a. Dissolve 1.3 g EDTA in 30 mL distilled water. Adjust the pH to 8.0 using 6 M NaOH. Add an additional 1 g EDTA, mix until dissolved, and re-adjust the pH to 8.0. Continue adding 1 g EDTA at a time, adjusting the pH back to 8.0 after each addition, until a total of 9.3 g EDTA has been added. Bring the final volume to 50 mL with distilled water. Store at 4 °C.

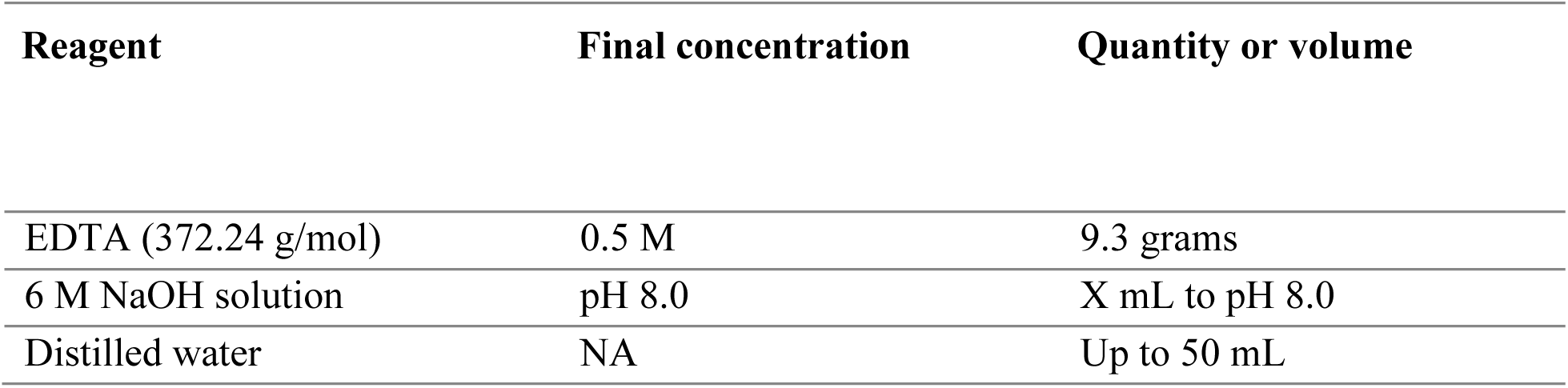

| Reagent | Final concentration | Quantity or volume |
| --- | --- | --- |
| EDTA (372.24 g/mol) | 0.5 M | 9.3 grams |
| 6 M NaOH solution | pH 8.0 | X mL to pH 8.0 |
| Distilled water | NA | Up to 50 mL |

16. 100x Protease inhibitor tablet solution (100 µL)

a. Store at -20 °C unless using the day of.

16. 100x Protease inhibitor tablet solution (100 μL) - a. Store at -20 °C unless using the day of.
| Reagent | Final concentration | Quantity or volume |
| --- | --- | --- |
| cOmplete Protease Inhibitor Cocktail | 100x | 1 tablet |
| Distilled water | NA | 500 μL |

17. 0.5 mg/mL Mass Spec Protease solution (40 µL)

a. Store at −20 °C. If the protease mix is used within one hour, it may be kept on ice instead of freezing.

17. 0.5 mg/mL Mass Spec Protease solution (40 μL) - a. Store at -20 °C. If the protease mix is used within one hour, it may be kept on ice instead of freezing.
| Reagent | Final concentration | Quantity or volume |
| --- | --- | --- |
| Commercially purchased MS-grade proteases | 0.5 μg/μL | 20 μg |
| 1 M Tris pH 7.8 solution | NA | 40 μL |

18. 1 M DTT solution (100 µL)

a. Prepare the solution fresh on the day of use and keep on ice until needed.

18. 1 M DTT solution (100 μL) - a. Prepare the solution fresh on the day of use and keep on ice until needed.
| Reagent | Final concentration | Quantity or volume |
| --- | --- | --- |
| DTT (154.25 g/mol) | 1 M | 15.42 mg |
| HPLC-grade water | NA | 100 μL |

19. 500 mM IAA solution (100 µL)

a. Prepare the solution fresh immediately before use, keep on ice, and protect it from light by keeping the tube covered or wrapped in foil when not in use.

| Reagent | Final concentration | Quantity or volume |
| --- | --- | --- |
| IAA (184.96 g/mol) | 500 mM | 9.2 mg |
| HPLC-grade water | NA | 100 μL |

20. 2.4 mg/mL PDET-1 solution (8.3 mL)

a. Store at 4 °C. Check pH with pH indicator strips 5.0-10.0

20. 2.4 mg/mL PDET-1 solution (8.3 mL) - a. Store at 4 °C. Check pH with pH indicator strips 5.0-10.0
| Reagent | Final concentration | Quantity or volume |
| --- | --- | --- |
| Commercially purchased PDET-1 | 2.4 mg/mL | 20 mg |
| 1 M Tris pH 7.8 solution | pH 7.5 | X mL |
| Distilled water | NA | To 8.3 mL |

21. 5 mg/mL HD-43 solution (4 ml)

a. Store at 4 °C. Check pH with pH indicator strips 5.0-10.0

21. 5 mg/mL HD-43 solution (4 ml) - a. Store at 4 °C. Check pH with pH indicator strips 5.0-10.0
| Reagent | Final concentration | Quantity or volume |
| --- | --- | --- |
| Commercially purchased HD-43 | 5 mg/mL | 20 mg |
| 1 M Tris pH 7.8 solution | pH 7.5 | X mL |
| Distilled water | NA | To 4 mL |

22. Solubilization buffer (5 mL)

a. Make fresh and keep on ice for use the day of.

22. Solubilization buffer (5 mL) - a. Make fresh and keep on ice for use the day of.
| Reagent | Final concentration | Quantity or volume |
| --- | --- | --- |
| 2.4 mg/mL PDET-1 solution | 2 mg/mL | 4.17 mL |
| 1 M Tris pH 7.8 solution | 50 mM | 250 μL |
| 1 M NaCl solution | 100 mM | 500 μL |
| 0.5 M EDTA pH 8.0 solution | 1 mM | 10 μL |
| 100x Protease Inhibitors solution | 1x | 50 μL |

23. C18 Resin Slurry (600 µL)

a. Make just before use.

23. C18 Resin Slurry (600 μL) - a. Make just before use.
| Reagent | Final concentration | Quantity or volume |
| --- | --- | --- |
| Commercially purchased C18 resin | NA | 200 μL of resin |
| Methanol | NA | 400 μL |

24. 10% Formic acid solution (10 mL)

a. Store at 4 °C.

24. 10% Formic acid solution (10 mL) - a. Store at 4 °C.
| Reagent | Final concentration | Quantity or volume |
| --- | --- | --- |
| Commercially purchased 100% Formic acid solution | 10% | 1 mL |
| HPLC-grade water | NA | 9 mL |

25. 0.1% Formic acid solution (10 mL)

a. Store at 4 °C.

25. 0.1% Formic acid solution (10 mL) a. Store at 4 °C.
| Reagent | Final concentration | Quantity or volume |
| --- | --- | --- |
| 10% Formic acid solution | 0.1% | 100 µL |
| HPLC-grade water | NA | 9.9 mL |

26. 40% Acetonitrile, 0.1% Formic acid solution (10 mL)

a. Store at 4 °C.

26. 40% Acetonitrile, 0.1% Formic acid solution (10 mL) a. Store at 4 °C.
| Reagent | Final concentration | Quantity or volume |
| --- | --- | --- |
| Commercially purchased 100% Acetonitrile solution | 40% | 4 mL |
| 10% Formic acid solution | 0.1% | 100 µL |
| HPLC-grade water | NA | 5.9 mL |

27. 1X Bradford Assay solution (50 mL)

a. Store at 4 °C.

27. 1X Bradford Assay solution (50 mL) a. Store at 4 °C.
| Reagent | Final concentration | Quantity or volume |
| --- | --- | --- |
| Commercially purchased Protein Assay Dye Reagent Concentrate | 1 X | 10 mL |
| Distilled water | 40 mL | 40 mL |

### Equipment

● Burner; Fisher Scientific; Catalogue number 01257557.
● Laboratory Lighter; Fisher Scientific; Catalogue number NC169758815.
● Pipette Controller; Fisher Scientific; Catalogue number 10-320-173.
● 10 mL polystyrene serological pipets; Fisher Scientific; Catalogue number 1367518.
● 7 and 15 mL Glass Douncer; Fisher Scientific; Catalogue number K885300007 and K8853000015.
● Eppendorf Table Top Centrifuge 5415D; Sigma-Aldrich; Catalogue number Z604062.
● Tabletop Ultracentrifuge Optima Max 130,000 RPM; Beckman Coulter.
● TLA-55 Fixed-Angle Rotor; Beckman Coulter; Catalogue number 366725.
● TLA-55 1.5 mL Polypropylene Tube with Snap-On Cap. Beckman Coulter; Product number 343169.
● C24 Incubator shaker; New Brunswick Scientific.
● C25 Incubator shaker; New Brunswick Scientific.
● M110L Microfluidizer Processor; Microfluidics Corporation.
● Labquake Rotisserie; ThermoFisher Scientific.
● Corning 125 mL and 1 L Erlenmeyer flasks; Fisher Scientific; Catalogue number 100414A and D.
● 1.5 mL Microcentrifuge Tubes; Fisher Scientific; Catalogue number 05408129.
● Nitrile Gloves; FroggaBio; Catalogue number FG100M-10.
● Bulk p1000, p200, and p10 pipette tips; Sarstedt; Catalogue number 70.1187.102, 70.3030.020, 70.3010.205.
● Scoopula; Fisher Scientific; Catalogue number 01257567.
● Magnetic Stir bars; Fisher Scientific; Catalogue number 1451357SIX.
● 100 mL, 250 mL, 500 mL, and 1L Media storage bottles; Fisher Scientific; Catalogue number FB800100, FB800250, FB800500, FB8001000.
● pH indicator strips 2.5-4.5; Millipore-Sigma; Catalogue number 1095410001.
● pH indicator strips 5.0-10.0; Millipore-Sigma; Catalogue number 1095330001.
● Ice; Provided by the university.
● Avanti J-E Centrifuge; Beckman Coulter.
● JA-25.50 Fixed-Angle Rotor-Aluminum; Beckman Coulter; Catalogue number 363055.
● JA 25.50 50 mL Polypropylene Bottle with Cap Assembly; Beckman Coulter; Catalogue number 361694.
● JLA10.500 fixed-angle aluminum rotor; Beckman Coulter; Catalogue number 369681.
● JLA10.500 500 mL polycarbonate bottles; Beckman Coulter; Catalogue number 369681.
● Genesys 10 UV-Vis Spectrophotometer; Fisher Scientific; Catalogue number 14385400.
● Falcon 50 mL Conical Centrifuge Tube; Fisher Scientific; Catalogue number 1443222.
● Falcon 15 mL Conical Centrifuge Tube; Fisher Scientific; Catalogue number 1495953B.
● Nutator; Clay Adams.
● Analog Vortex Mixer; Fisher Scientific; Catalogue number 02215414.
● Optima LE-80k Ultracentrifuge; Beckman Coulter.
● TI-45 Fixed-Angle Titanium Rotor; Beckman Coulter; Catalogue number 339160.
● TI-45 Round Top Ultra-Clear Tube; Beckman Coulter; Catalogue number 345778.
● Analytical Scale TL-64; Denver Instruments Company.
● Toploading Scale APX-2001; Denver Instruments Company.
● Hot Plate Stirrer PC-351; Corning.
● pH meter AB15; Fisherbrand accumet.
● Needle; Becton Dickinson; Catalogue number 305196.
● Standard-Duty Vacuum Pump; Fisher Scientific; Model number 2545B-01.
● Thermo Savant Speed Vac Concentrator SC110; Marshall Scientific.
● Savant Speed Vac Refrigerated Vapor Trap; Fisher Scientific; Catalogue number 13875349.
● Gel-Loading Tips; Fisher Scientific; Catalogue number 02707181.
● Semi-micro cuvette; Millipore-Sigma; Catalogue number BR759015-100EA.
● 4 °C fridge, -20 °C, and -80 °C freezer.

### Procedure

#### A. *E. coli* Culture Growth and Cell Harvesting

1. Activate the CO₂-connected Bunsen burner by turning the valve to release gas and ignite the burner using a lighter.
2. Near a lit Bunsen burner (to reduce the risk of contamination from airborne particles), add 50 mL of LB medium to a sterile 125 mL Erlenmeyer flask.
3. Remove the BL21 *E. coli* glycerol stock from the −80 °C freezer. Using a sterile P200 pipette tip, scrape a small frozen portion of the glycerol stock.
4. Drop the frozen scrape of the glycerol stock into the LB medium in the 125 mL Erlenmeyer flask while working near the Bunsen burner flame.
5. Turn off the Bunsen burner and incubate the Erlenmeyer flask containing the bacterial scrape in the C24 incubator shaker at 37 °C and 200 rpm for 12–16 hours to generate the bacterial starter culture.

➢ **Caution: The LB medium does not contain antibiotics, so any contaminants introduced during handling will also grow in the culture. Perform all steps near an active Bunsen burner to maintain sterility.**
6. The next day, ignite the Bunsen burner using a striker. Working near the flame, transfer LB medium into a 1 L Erlenmeyer flask and use the flask graduations to measure 1 L, as the medium cannot be transferred to a graduated cylinder due to the absence of antibiotics.
7. Remove the bacterial starter culture and, working near the flame, transfer 10 mL into the 1 L Erlenmeyer flask containing 1 L of fresh LB medium. Cover the mouth of the flask with aluminum foil to prevent contaminants from entering the culture during incubation.
8. Transfer the flask to the C25 incubator shaker and grow the culture at 37 °C with shaking at 200 rpm for 8–16 hours.
9. After incubation, remove the bacterial culture from the C25 shaker. Transfer 500 mL of the culture into each of two JLA 10.500 polypropylene centrifuge tubes. Weigh the tubes to ensure they are balanced within 5 grams of each other.
10. Place the JLA-10.500 rotor into the Avanti J-E centrifuge.
11. Place the two centrifuge tubes into JLA-10.500 biosafety liquid canisters and position them opposite each other in the JLA-10.500 rotor. Fill the remaining rotor positions with empty biosafety liquid canisters to ensure the rotor is fully balanced.

➢ **Critical: All slots in the JLA-10.500 rotor must be filled before centrifugation. Any unused positions must be occupied with empty biosafety liquid canisters to ensure proper rotor balance during the spin.**
12. Centrifuge at 6,000 × g for 15 minutes.
13. After centrifugation, discard the supernatant (spent LB media) down the drain, leaving the bacterial pellet in the tube.
14. Resuspend the bacterial pellet in 10 mL of TSG buffer per liter of LB culture used to generate the pellet. Add 10 mL of TSG buffer to one of the 500 mL centrifuge bottles containing the pellet and homogenize it using the glass pestle from the 15 mL glass douncer set in combination with vortexing until the pellet is fully resuspended.
15. After the pellet has been dislodged and homogenized, transfer the 10 mL suspension using a serological pipette to the second 500 mL centrifuge bottle containing the remaining bacterial pellet. Homogenize again using the glass pestle and vortexing until the pellet is fully resuspended.
16. Once both pellets are dislodged, transfer the homogenate to the 15 mL glass douncer by pouring it from the centrifuge bottle. If necessary, add 1–2 mL of TSG buffer to the centrifuge bottle to resuspend any remaining pellet fragments and transfer this rinse to the douncer as well.
17. Dounce the homogenate in the 15 mL glass douncer until all visible clumps are dispersed and the sample appears fully homogeneous.
18. Transfer the homogenate to a 50 mL Falcon tube by pouring it out of the glass douncer. If any homogenate remains in the douncer, add 1–2 mL of buffer to rinse it, then transfer the rinse into the same 50 mL Falcon tube.
19. Store the homogenate at −20 °C until further use.

### B. Bacterial Homogenate Lysis

1. Remove the bacterial homogenate from −20 °C and thaw it on ice for approximately 30 minutes.

i. If pressed for time, the sample can be thawed in a beaker of room-temperature water for 5–10 minutes.
2. Prepare the M110L microfluidizer (stored in 20% ethanol) for use:

i. Insert the glass funnel to the machine inlet.
ii. Pack the bed of the microfluidizer with ice to keep the sample cold during processing.
3. Open the valve on top of the compressed air gas cylinder while keeping the microfluidizer air switch switch closed.
4. Set the microfluidizer pressure to 12,000 psi by turning the pressure valve clockwise.
5. Flush the machine with distilled water to remove the residual 20% ethanol used for storage.

i. Turn the microfluidizer air intake switch to start the machine and allow the sample to flow from the glass funnel into the microfluidizer.
ii. A “click” corresponds to the sound produced when a portion of the solution in the intake valve is pushed through the machine in a stepwise manner.
iii. One wash corresponds to three clicks from the machine.
iv. Perform three washes with distilled water.
v. After washing, discard the excess water from the glass funnel.

➢ **Critical: Ensure that no air enters the machine, as air bubbles can clog the system. Always ensure that liquid is present in the glass intake funnel before turning the machine on.**
6. Add the thawed bacterial homogenate to the intake funnel and pass it through the microfluidizer five times.

➢ **Critical: Collect the output in a 50 mL Falcon tube by placing the outlet tubing into the tube. Otherwise, the sample will be directed to waste and lost.**
7. After the fifth pass, add approximately 5 mL of TSG buffer to the glass intake funnel. Turn the machine on to flush any remaining homogenate from the tubing, and collect this with the rest of the lysate. Perform again until you the tubing is clear of homogenate.
8. Keep the lysed homogenate on ice.
9. Clean the machine immediately after use:

i. Perform three washes with distilled water.
ii. Follow with three washes using 20% ethanol.
iii. Leave the machine filled with 20% ethanol for storage to prevent bacterial growth in the tubing.
iv. Remove the glass funnel and top off the intake with 20% ethanol. Place a bottle cap over the intake to prevent the 20% ethanol from drying out

**➢ Critical: If the machine runs dry, it will become unusable and require the pressure cell to be cleaned to restore proper operation.**
v. Close the valve on top of the compressed air gas cylinder.

**Figure 1.**
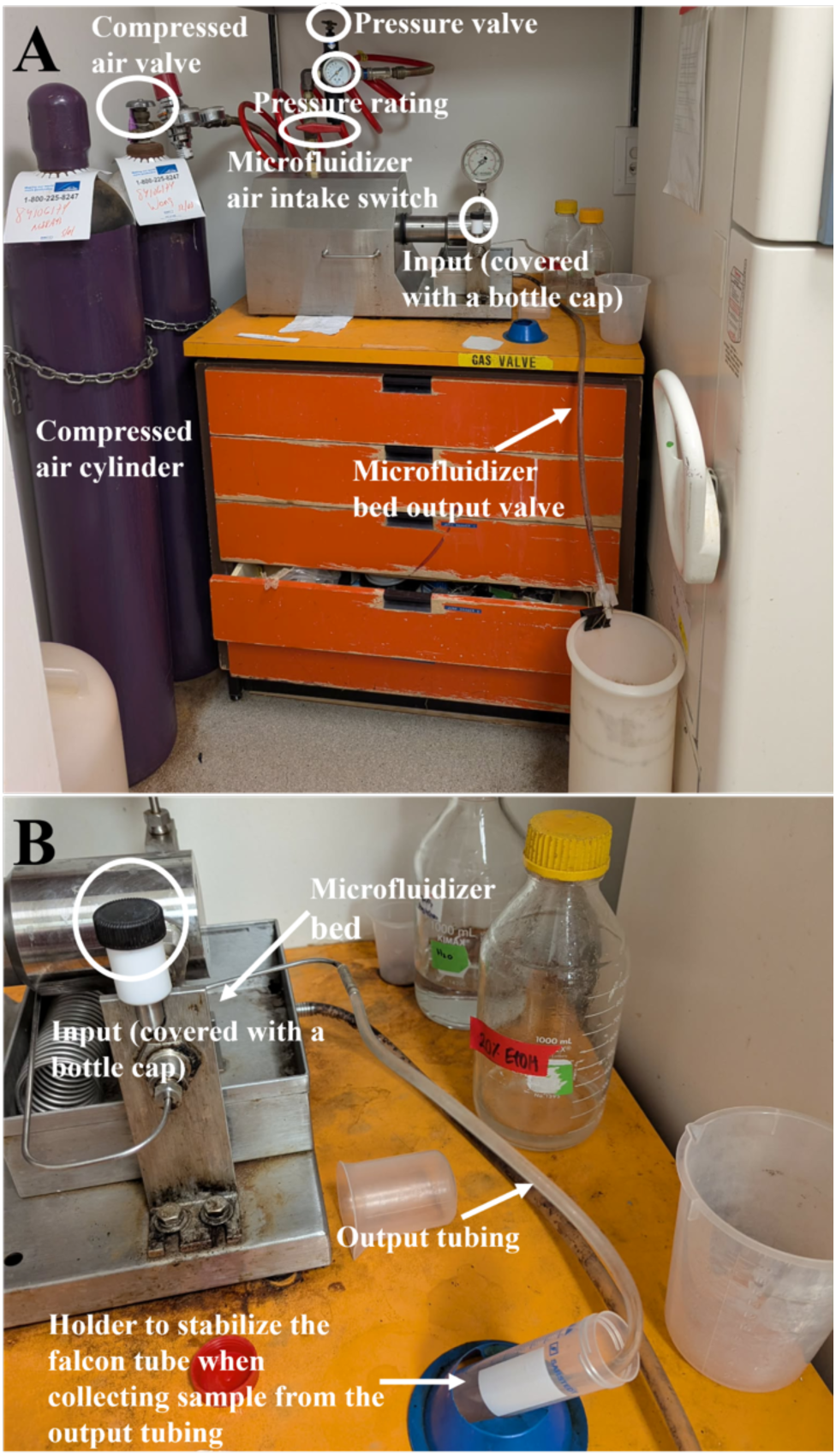
Visual overview of the microfluidizer used for bacterial cell lysis.

#### C. Isolation of Crude Membrane Fraction from *E. coli*

1. Transfer the homogenate into two JA-25.50 50 mL polypropylene centrifuge bottles, topping off with TSG buffer to balance the tubes. Ensure the tubes are within 1 g of each other.
2. Centrifuge using a JA-25.50 rotor in the Avanti J-E centrifuge at 4,350 × g for 10 minutes to pellet the unlysed cells.
3. After centrifugation, the supernatant will contain the bacterial lysate and the pellet will contain unlysed cells. Carefully remove the supernatant while avoiding disturbance of the pellet. It is acceptable to leave a small amount of supernatant behind to ensure that no unlysed cells are collected.
4. Add an equal volume of bleach to the unlysed cell pellet and allow it to sit at room temperature for 30 minutes for decontamination. Afterward, dispose of the mixture down the drain and flush with running water for 15 minutes.
5. Transfer the bacterial lysate into TI-45 ultra-clear round top centrifuge bottles, distributing the lysate evenly between two bottles. Top up each bottle with TSG buffer to the maximum fill volume to ensure proper balance.

➢ **Caution – After topping off the bottle, ensure that no more than one air bubble remains inside the tube. If multiple bubbles are present, reopen the bottle and top off again, placing the cap on carefully to minimize bubble formation. Check for bubbles by gently inverting the bottle. Limiting air bubbles helps prevent leakage during the spin, which can lead to sample loss and rotor imbalance that will stop the centrifuge.**
6. Load the tubes into the TI-45 rotor. The rotor should be stored in the cold room when not in use. Place the rotor into the Optima LE-80K ultracentrifuge.
7. After placing the rotor in the centrifuge, close the lid and allow the vacuum to build. Once the vacuum reaches 750, start the run at 40,000 rpm for 45 minutes (corresponding to 125,440 × g, calculated using the rotor calculator on the Beckman Coulter website).
8. After centrifugation, remove the tubes and place them on ice. Return the TI-45 rotor to the cold room for storage.

➢ **Caution – Inspect and clean the ultracentrifuge and rotor for any lysate that may have leaked during the spin. Failure to do so may lead to bacterial or fungal growth inside the ultracentrifuge or rotor.**
9. The supernatant contains the soluble fraction of the *E. coli* lysate, while the pellet contains the membrane fraction (crude membrane).

i. Remove the supernatant and store it at −20 °C.
10. Resuspend the crude membrane pellet in TSG buffer using 5 mL of TSG per liter of LB culture used to generate the crude membrane. This corresponds to 2.5 mL per centrifuge bottle.

i. Add the TSG buffer, then dislodge the pellet using a P1000 pipette tip. After the pellet has been loosened, cut the end of a new P1000 pipette tip to widen the opening and pipette up and down to break up the remaining chunks until they are small enough to be easily aspirated.
11. Transfer the resuspended crude membrane to a 7 mL glass douncer.

i. Add 500 µL of TSG to each centrifuge bottle to collect any remaining chunks of crude membrane. Cut the end off a P1000 pipette tip and use it to transfer all remaining material to the 7 mL glass douncer.
12. Homogenize the sample by douncing until the crude membrane suspension is uniform and no visible chunks remain.
13. Transfer the crude membrane suspension to a 15 mL Falcon tube.
14. Determine the protein concentration using the Bradford assay.

i. Add 1 mL of 1× Bradford assay solution to four semi-micro cuvettes.
ii. Add 2 µL of TSG to the first cuvette (blank). Add 1 µL of crude membrane to the second cuvette, 2 µL to the third cuvette, and 3 µL to the fourth cuvette.
iii. Mix each cuvette by pipetting the solution up and down with a P1000, using a fresh tip for each cuvette.
iv. Place the four cuvettes into the cuvette holder of the Genesys 10 UV–Vis spectrophotometer as follows: the TSG blank in slot B, 1 µL crude membrane in slot 1, 2 µL crude membrane in slot 2, and 3 µL crude membrane in slot 3.
v. On the instrument, press the Test button. Then press the down arrow once to highlight “Standard Curve.”
vi. Name the test “Bradford.” Set the wavelength to “595 nm”, Ref. Wavelength Correction to “Off”, and Curve Fit to “Linear Through Zero.’ Set the number of standards to “Five”, units to “mg/mL”, sample positioner to “Auto 6”, and the number of samples to “five”.
vii. Run the test.
viii. Calculate the average protein concentration across the three cuvettes. Divide the value from slot 2 by 2 and the value from slot 3 by 3 to account for the 2 µL and 3 µL sample volumes added, respectively, then average the normalized values with the value from slot 1.

**➢ The instrument has a maximum measurable concentration of 25 mg/mL. Any reading ≥20 mg/mL should be considered outside the reliable measurement range and excluded when calculating the average protein concentration.**
**➢ The Bradford assay provides an estimate of the protein concentration in the crude membrane and can be used to guide subsequent experiments, but it should not be considered an exact measurement. For assays requiring highly accurate protein quantification, protein levels should be further normalized by running and quantifying samples on SDS–PAGE gels to account for potential protein contamination.**
15. Record the protein concentration on the Falcon tube containing the crude membrane and store it at −20 °C until further use.

➢ **Caution - The crude membrane is stable at −20 °C for extended periods (up to ∼1 year). However, protein yield decreases with each freeze–thaw cycle. It is recommended to repeat the Bradford assay after each freeze–thaw cycle to determine the updated protein concentration.**

#### D. Solubilization of the Membrane Proteome Using Peptergent (PDET-1)

1. Remove the crude membrane from −20 °C and allow it to thaw on ice for approximately 30 minutes.

a. If needed, the crude membrane can be thawed more quickly by placing the tube in a beaker of room-temperature water for 5–10 minutes.
b. While the crude membrane is thawing, prepare the solubilization buffer.
2. Resuspend 1 mg of crude membrane protein in 500 µL of solubilization buffer (1 mg crude membrane to 1 mg PDET-1).

a. Adjust the crude membrane concentration to 2 mg/mL prior to solubilization so that 500 µL of crude membrane can be mixed with 500 µL of solubilization buffer, resulting in a final volume of 1 mL. If the crude membrane preparation is more dilute than 2 mg/mL, still use 500 µL of the membrane suspension so that the final volume remains 1 mL.
b. Set aside 10 µL of the 2 mg/mL crude membrane for troubleshooting by SDS–PAGE analysis.
3. Incubate the suspension on a rotisserie shaker in the cold room for 90 minutes to allow membrane protein extraction with PDET-1.
4. After the 90-minute incubation, transfer 0.5 mL of the suspension into two separate TLA-55 1.5 mL polypropylene tubes.

a. Visually check that the samples have equal volumes for proper balancing. If one tube has a lower volume, add TS buffer to match the volume of the other tube.
5. Place the tubes into a TLA-55 fixed-angle rotor. The rotor should be stored in the cold room when not in use.
6. Place the rotor in the Optima Max tabletop ultracentrifuge, ensuring the center of the rotor clicks securely into place before starting the spin.
7. Centrifuge at 55,000 rpm for 15 minutes (corresponding to 135,520 × g, calculated using the rotor calculator on the Beckman Coulter website).
8. The supernatant contains the clarified membrane proteome extract solubilized with PDET-1, while the pellet contains the insoluble material.

a. Set aside the pellet for SDS–PAGE troubleshooting in case your protein of interest is not successfully extracted.
9. Transfer the supernatant to a new microcentrifuge tube.

a. Set aside 10 µL of the extract for SDS–PAGE troubleshooting.

#### E. Peptide Exchange from PepTergent to His-Tagged Peptidisc

1. Take 500 µL of the PDET-1 extract and add 200 µL of 5 mg/mL HD-43 solution, resulting in a 1:1 ratio of PDET-1 to HD-43.

a. The extract volume may vary after clarification depending on the size of the insoluble pellet or on the introduction of bubbles during centrifugation due to tube imbalance. For this step, maintain a 500 µL PDET-1 extract-to-200 µL HD-43 ratio, scaling the volumes as needed to preserve the 1:1 PDET-1-to-HD-43 ratio during the exchange.
2. Incubate the mixture on a rotisserie shaker in the cold room for 10 minutes to allow peptide exchange.

#### F. Affinity Purification Enabling Exchange of PDET-1 into HD-43 and the Enrichment of Reconstituted Membrane Proteins

1. Prepare the Ni-NTA resin during the exchange step. The resin is stored as a 50:50 slurry of resin in 30% ethanol. Pipette 200 µL of the Ni-NTA slurry into a microcentrifuge tube using a cut P200 pipette tip to prevent the slurry from sticking to the tip.

a. Centrifuge at 2,000 × g for 2 minutes.
b. Carefully remove the 30% ethanol supernatant, ensuring that the resin is not disturbed.

➢ **Caution – The resin may be aspirated when removing liquid near the pellet. If this occurs, centrifuge the tube again at 2,000 × g for 2 minutes to re-pellet the resin. If additional liquid cannot be removed without disturbing the resin, leave the remaining volume in the tube and proceed to the next step.**
c. Resuspend the resin in 500 µL of distilled water and incubate for 5 minutes at room temperature on the nutator.
d. Centrifuge again at 2,000 × g for 2 minutes, then aspirate the water.
e. Resuspend the resin in 500 µL of TS buffer and keep it at room temperature on the nutator until further use.
2. Pellet the Ni-NTA resin by centrifugation at 2,000 × g for 2 minutes, then remove the TS buffer.
3. Add the 1:1 PDET-1:HD-43 extract to the Ni-NTA resin and incubate for 1 hour on a rotisserie rotator in the cold room to allow binding of His-tagged Peptidisc reconstituted MPs to the Ni-NTA resin.

➢ **Critical – Binding of the His-tagged Peptidisc to the Ni-NTA resin enables the selective capture of successfully exchanged and reconstituted membrane proteins in HD-43, separating them from contaminating soluble proteins and MPs whose peptide was not successfully exchanged for HD-43.**
➢ **Caution – A total volume of 500 µL–1 mL is recommended during incubation to allow proper mixing on the rotisserie so that the liquid moves freely with the resin. If the volume is insufficient, add TS buffer to bring the total volume to 500 µL to enable proper mixing.**
➢ **Pause point – This step can be extended from 1 hour to overnight if needed. Although the protocol specifies 1 hour, this step can serve as a convenient stopping point for researchers.**
4. After incubation, centrifuge at 2,000 × g for 2 minutes to pellet the resin.
5. Carefully remove the flow-through (unbound proteins) and store it at 4 °C for SDS–PAGE analysis.
6. Wash the bound Ni-NTA resin three times with TS buffer.

a. Add 1 mL of TS buffer and rotate for 5 minutes on the rotisserie in the cold room.
b. Centrifuge at 2,000 × g for 2 minutes, then discard the wash.
7. After washing the Ni-NTA resin and removing the TS buffer, elute the bound proteins with 200 µL of TS buffer with 600 mM imidazole solution for 20 minutes on the rotisserie in the cold room.

➢ **Pause point – Similar to the PDET-1/HD-43 extract binding step to the Ni-NTA resin, this step can also be extended overnight, serving as a pause point for researchers.**
8. Centrifuge at 2,000 × g for 2 minutes, then carefully collect the eluate, ensuring that no resin is carried over.
9. Perform an additional centrifugation if necessary to recover any remaining eluate.

a. **Caution – Multiple spins are often required to safely remove the eluate without disturbing the resin.**
10. Transfer the eluate to a new microcentrifuge tube.

a. Set aside 10 µL for SDS–PAGE troubleshooting analysis.
11. Determine the protein concentration using the Bradford assay.

a. Use 3 µL TS buffer containing 600 mM imidazole as the blank. For the samples, add 1 µL, 3 µL, and 5 µL of the eluate to 1 mL of 1× Bradford assay solution. Average the protein concentrations obtained from these three measurements. The methodology for performing the Bradford assay is described in Section C, Step 14.
12. After determining the protein concentration, remove a volume corresponding to 100 µg of protein. Dilute this to a final volume of 100 µL using TS buffer, resulting in a 100 µL solution containing 100 µg of the eluted Peptidisc-reconstituted membrane proteome. Keep this solution on ice.

a. If the sample is too dilute to obtain 100 µg in 100 µL, use the volume corresponding to 100 µg of protein and proceed with the MS preparation steps using the adjusted volume.
13. Store the remaining eluted membrane proteome at −20 °C.

➢ **Caution – The Peptidisc-reconstituted membrane proteome can be stored for up to one week with little to no protein loss. If the sample will be used within a week, store it at 4 °C. For longer storage, keep it at −20 °C. Protein degradation can occur with freeze–thaw cycles, so the protein concentration should be re-measured by Bradford assay after each freeze–thaw cycle.**
14. Before proceeding with MS sample preparation, run the samples on SDS–PAGE to verify that the preparation was successful (example in Figure 2). If any unexpected results are observed, it is recommended to pause the protocol and troubleshoot the issue before continuing. Do not proceed to MS analysis until the problem has been resolved.

**➢ Critical – Based on the protocol above, run the following samples on SDS–PAGE: crude membrane, PDET-1 extract, PDET-1 insoluble pellet (only if the protein of interest is not being extracted), Ni-NTA flow-through, and Ni-NTA elution. We recommend loading 5 µL of each sample onto the gel to prevent overloading.**

**Figure 2:**
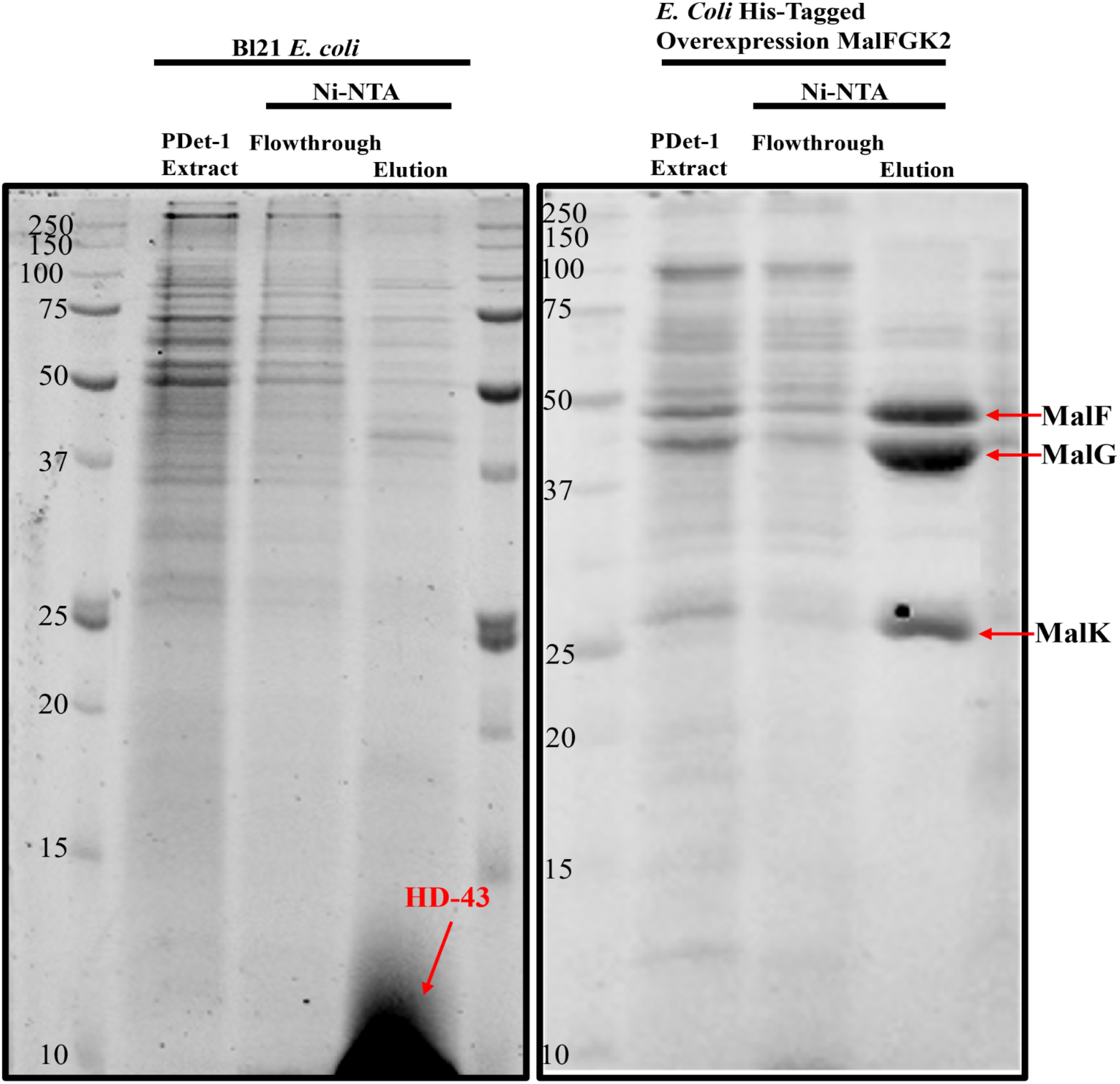
PDET-1 extraction and purification of the membrane proteome and the His-tagged MalFGK_2_ complex. **Left**: SDS–PAGE analysis of the membrane proteome obtained with PDET-1 + HD43 before (extract) and after (eluate) Ni-NTA purification. **Right**: SDS–PAGE analysis of the His-tagged MalFGK_2_ solubilized with PDET-1, before (extract) and after (eluate) Ni-NTA-based purification. The flow-through and molecular weight markers are also presented.

**Figure 3:**
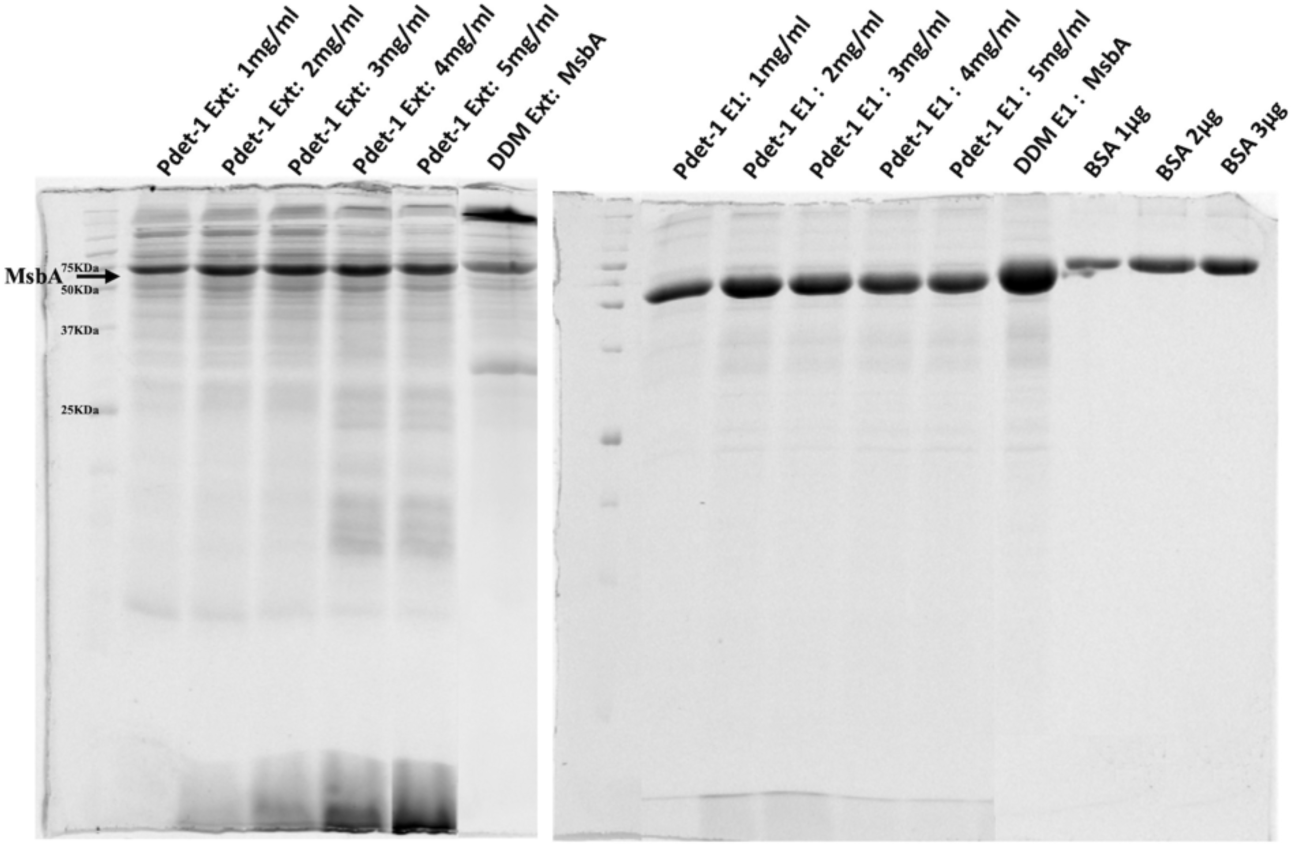
Peptergent-based purification of the ABC transporter his-MsbA. The membrane preparation enriched for MsbA (1 mg) was incubated with PDET-1 at the indicated concentration before ultra-centrifugation. **Left**: An aliquot of the obtained supernatant (the Peptergent extract) was analyzed by SDS–PAGE and Coomassie-blue staining. The remaining Peptergent extract was incubated with Ni-NTA beads. **Right**: An aliquot of the eluate was analyzed by SDS–PAGE and Coomassie-blue staining of the gel. For reference, an aliquot of the DDM (dodecyl maltoside) detergent extract before and after Ni-NTA-based purification is loaded on the gels. The low-molecular-weight band seen at the bottom of the gel is the PDET-1 peptide.

**Figure 4.**
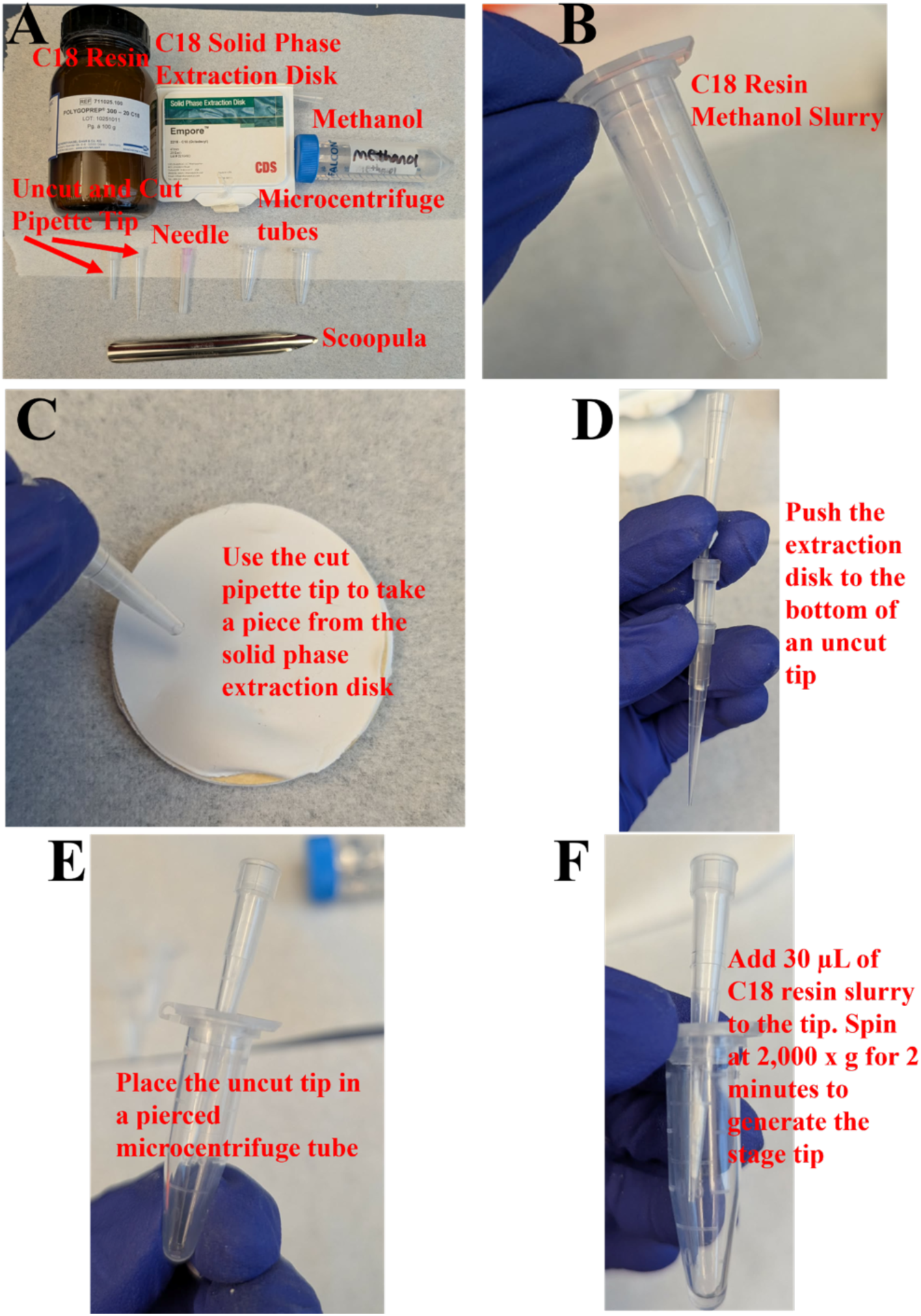
Visual workflow for preparing custom StageTips.

#### G. Bottom-Up MS Sample Preparation

1. Add 36.7 mg of urea to 100 µL containing 100 µg of the eluted membrane proteome sample to reach a final concentration of 6 M urea. Vortex to mix.
2. Incubate at room temperature on the nutator for 30 minutes.
3. Add 1 µL of 1 M DTT to the sample to achieve a final concentration of 10 mM. Incubate on the nutator at room temperature for 1 hour.
4. Add 4 µL of 500 mM IAA to the sample to achieve a final concentration of 20 mM, and incubate at room temperature in the dark on the nutator for 30 minutes.
5. Add 1 µL of 1 M DTT to the sample to reach a final concentration of 20 mM. Incubate at room temperature on the nutator for 30 minutes.
6. Add 500 µL of 50 mM ammonium bicarbonate to dilute the urea concentration to 1 M.

a. High urea concentrations disrupt the activity of MS-grade proteases.
7. Add 2 µL of 0.5 µg/µL protease solution (1:100 trypsin-to-sample ratio) and incubate overnight at room temperature on the nutator.
8. The following day, add an additional 2 µL of 0.5 µg/µL protease solution to reach a final enzyme-to-protein ratio of 1:50, and incubate overnight at room temperature on the nutator.

➢ **Pause point – Either of the trypsin digestion steps can be extended to incubate over the weekend, allowing researchers to pause the protocol and resume on the next working day.**

#### H. C18 StageTip Peptide Desalting and Drying

1. Prepare the C18 resin slurry (Figure 3B), 0.1% formic acid solution, 10% formic acid solution, and 40% acetonitrile in 0.1% formic acid solution.
2. Use scissors to cut the end off a 200 µL pipette tip. Using the bottom of the cut tip, take a piece of the C18 solid phase extraction disk (Figure 3C) and transfer it into a new, uncut 200 µL pipette tip. Use a gel loading tip to push the extraction disk to the bottom of the uncut tip (Figure 3D).
3. Using a needle, punch a hole in the lid of a new microcentrifuge tube and insert the tip with the extraction disk (Figure 3E).
4. Pipette 30 µL of the C18 resin slurry into each prepared StageTip placed in its respective pierced microcentrifuge tube. Then centrifuge at 2,000 × g for 2 minutes generating the stage tip (Figure 3F).
5. After the spin, add 75 µL of methanol and centrifuge at 2,000 × g for 2 minutes.
6. Wash the StageTips twice with 100 µL of 0.1% formic acid solution, centrifuging at 2,000 × g for 2 minutes after each wash.
7. To the trypsin-digested samples (now 600 µL), add 20 µL of 10% formic acid solution. Check the pH of the sample and add additional formic acid if necessary to reach pH 3. Use a pH indicator strip 2.5-4.5 to check the pH.
8. Remove the StageTip from the microcentrifuge tube and pour the flow-through from the bottom of the tube into a waste container. Then return the StageTip to its corresponding tube.

**a. Critical - Ensure the labels on the StageTip microcentrifuge tubes are clearly visible and readable, as mixing up samples at this stage could compromise over a week of work.**
9. Add 100 µL of the pH 3 trypsin-digested sample to the corresponding C18 StageTip and centrifuge at 2,000 × g for 2 minutes. Repeat this process, adding 100 µL of the digested sample per cycle, until the entire sample has been loaded onto the StageTip.

a. Discard the flow-through after each spin to prevent it from contacting the StageTip. If the flow-through level rises and begins to touch the StageTip, pour it out before continuing.
b. If the C18 resin becomes clogged, increase the centrifugal force to 3,000 × 2. g. If clogging persists, extend the spin time to 5 minutes.
c. In the worst-case scenario, use a syringe to gently force the liquid through the C18 StageTip.
10. Once the entire sample has been loaded onto the StageTip, wash the resin three times with 100 µL of 0.1% formic acid solution, centrifuging at 2,000 × g for 2 minutes for each wash.
11. After the final 0.1% formic acid wash, discard the flow-through and perform an additional dry spin at 2,000 x g for 2 minutes to ensure no liquid remains in the StageTip before elution.
12. Transfer the StageTip to a new pierced microcentrifuge tube.
13. Elute the bound peptides from the StageTip with 150 µL of 40% acetonitrile in 0.1% formic acid solution.

a. Perform this stepwise, eluting with 50 µL of 40% acetonitrile in 0.1% formic acid solution per step and centrifuging at 2,000 × g for 2 minutes each time, for a total of three spins.
14. After elution, transfer the entire eluate to a new microcentrifuge tube.
15. Place the sample in a lyophilizer/dryer connected to a vacuum pump and liquid condenser and dry on low heat until all liquid has evaporated (2–4 hours). Ensure the cap is open while in the machine to allow evaporation of the solvent.

➢ **Pause point – If needed, this step can be performed overnight. Ensure the heat is set to low if the sample is left to dry overnight.**
16. After the sample has dried, check the tube to ensure no liquid remains. If the sample is completely dry, seal the tube with Parafilm around the cap and store the samples for MS analysis.

### General notes and Troubleshooting

- This workflow is demonstrated here using *E. coli* BL21 crude membranes, but it is equally applicable to tissues and other cell types. The primary variables between sample types are the yield of crude membrane obtained from the starting material and the solubilization efficiency of the membrane proteome in PDET-1. Although not shown here, we have also successfully applied this method to mammalian cell lines and mouse organ tissues.
- It is recommended to analyze samples by SDS–PAGE at key steps of the workflow to monitor protein recovery and sample quality. For this protocol, these steps include the crude membrane, PDET-1-solubilized crude membrane extract, insoluble crude membrane fraction, Ni-NTA flow-through, and Ni-NTA eluent. Analyzing these samples helps determine whether the experiment proceeded correctly. One possible limitation is using insufficient crude membrane, which can result in inefficient solubilization and consequently weak band intensity on SDS–PAGE. Because PDET-1 and Peptidisc scaffolds are peptides, they will also be detected by the Bradford assay. It is therefore important to examine samples on SDS–PAGE to confirm that the preparation contains the membrane proteome, rather than the peptide scaffold alone, which could otherwise inflate the Bradford protein concentration measurement.
- If the protein of interest is not solubilizing, solubilize the crude membrane pellet with protein loading buffer and run the sample on an SDS–PAGE gel to determine whether the protein remains in the pellet. If the protein is still not solubilized, it may be necessary to use a detergent-based solubilization method.
- Other sample quantities can also be prepared for MS analysis. In this protocol, 100 µg of protein is processed. We recommend a minimum input of 50 µg.
- This is a long protocol, so take care when labeling samples at each step. If performing the workflow for the first time, we recommend starting with 1–3 samples to become familiar with the procedure. Once confident with the results, the protocol can be scaled to larger sample sets for full experiments. Careful labeling is especially critical during the StageTip step, as pipetting the wrong sample into the wrong StageTip can compromise over a week of work and lead to significant reagent and sample loss.
- Membrane proteins are stable in crude membrane preparations and in Peptidisc, so if a break is needed it is recommended to pause the protocol and resume the following day. Leaving samples overnight in the cold room will not appreciably affect the results. It is better to proceed carefully and obtain reliable, reproducible results rather than rushing through the protocol to complete it in a single session.

### Expected Outcomes

Application of the Peptergent-based workflow described here is expected to yield efficient extraction of membrane proteins directly from biological membranes while preserving their native organization. Following incubation with PDET-1, a substantial fraction of integral membrane proteins is solubilized into the supernatant after ultracentrifugation, while insoluble material is retained in the pellet. SDS–PAGE analysis typically reveals a broad distribution of proteins in the extract, including high–molecular–weight and multi-pass membrane proteins that are often underrepresented using detergent-based methods. Subsequent exchange into His-tagged Peptidiscs enables selective enrichment of reconstituted membrane proteins by Ni-NTA affinity purification. This step results in depletion of soluble contaminants and enrichment of membrane-associated species, as illustrated in Figures 2 and 3. Successful exchange and purification are typically evidenced by increased band intensity and reduced background in the elution fraction compared to the initial extract and flow-through.

As presented, downstream processing for bottom-up mass spectrometry yields peptide samples suitable for LC–MS/MS analysis without the need for detergent removal. In our experience, this workflow provides substantial coverage of the membrane proteome relative to conventional detergent-based extraction methods, including enhanced detection precision for hydrophobic and low-abundance proteins (8). The method is broadly applicable across sample types, including bacterial membranes, cultured mammalian cells, and tissue-derived fractions, and supports reproducible proteomic profiling of membrane protein composition and remodeling (22).

## Acknowledgments

Conceptualization, F.A. F.D.v.H.; Investigation, F.A. Writing—Original Draft, F.A. A.B.; Writing—Review & Editing, F.A. A.B. F.D.v.H.; Funding acquisition, A.B. F.D.v.H.; Supervision, F.D.v.H. The work is supported by an NSERC Discovery grant. A. B. is supported by a UBC 4-year fellowship, an Amplify Doctoral Award from Triangle (Training a new generation of researchers in gastroenterology and liver), and a Mast3 (Mass Spectrometry Team Training and Transition) scholarship.

